# CeGAL: revisiting a widespread fungal-specific TF family using an *in silico* error-aware approach to identify missing zinc cluster domains

**DOI:** 10.1101/2022.06.15.496365

**Authors:** Claudine Mayer, Arthur Vogt, Tuba Uslu, Nicolas Scalzitti, Olivier Poch, Julie D. Thompson

## Abstract

Transcription factors (TF) regulate gene activity in eukaryotic cells by binding specific regions of genomic DNA. In fungi, the most abundant TF class contains a fungal-specific ‘GAL4-like’ Zn2C6 DNA binding domain (DBD), while the second class contains another fungal-specific domain, known as ‘fungal_trans’ or Middle Homology Domain (MHD), whose function remains largely uncharacterized. Remarkably, almost a third of MHD-containing TF in public sequence databases apparently lack DNA binding activity, since they are not predicted to contain a DBD. Here, we reassess the domain organization of these ‘MHD-only’ proteins using an *in silico* error-aware approach. Our large-scale analysis of ~17000 MHD-only TF sequences showed that the vast majority (>90%) result from gene annotation errors, thus contradicting previous findings that the MHD-only TF are widespread in fungi. We show that they are in fact exceptional cases, and that the Zn2C6-MHD domain pair represents the canonical domain signature defining a new TF family composed of two fungal-specific domains. We call this family CeGAL, after the most characterized members: **Ce**p3, whose 3D structure has been determined and **GAL**4, an archetypal eukaryotic TF. This definition should improve the classification of the Zn2C6 TF and provide critical insights into fungal gene regulatory networks.

**IMPORTANCE:** In fungi, extensive efforts focus on genome-wide characterization of potential Transcription Factors (TFs) and their targets genes to provide a better understanding of fungal processes and a rational for transcriptional manipulation. The second most abundant families of fungal-specific TFs, characterized by a Middle Homology Domain, are major regulators of primary and secondary metabolisms, multidrug resistance and virulence. Remarkably, one third of these TFs do not have a DNA Binding Domain (DBD-orphan) and thus are excluded from genome-wide studies. This particularity has been the subject of debate for many years. By computationally inspecting the close genomic environment of about 20,000 DBD-orphan TFs from a wide range of fungal species, we reveal that more than 90% contained sequences encoding a zinc-finger DBD. This analysis implies that the arrays of DBD containing TFs and their control DNA-sequences in target genes need to be reconsidered and expands the combinatorial regulation degree of the crucial fungal processes controlled by this TF family.

## INTRODUCTION

Transcription factors (TF) are essential for the regulation of expression pathways in eukaryotes by binding genomic DNA *via* a DNA binding domain (DBD) for example composed of a zinc finger structural motif (1). The ‘classical’ zinc finger domain coordinates a single zinc atom with a combination of 4 amino acids, usually cysteine or histidine. However, the Zn2C6 domain (also called Zn(II)2Cys6, Zn2/Cys6 or Zn(2)-Cys(6) binuclear cluster domain) is an atypical zinc finger, where the well-conserved CX{2}CX{6}CX{5,16}CX{2}CX{6,8}C motif contains six conserved cysteines that coordinate two zinc atoms to establish correct folding of the zinc c1uster domain (2, 3). The Zn2C6 domain defines the GAL4-like Zn2C6-TF family, which is quasi-specific to fungi and observed ubiquitously in all fungal species, where it represents the most abundant TF family in each species (4–8). Zn2C6-TF are involved in a wide range of functions from primary and secondary metabolisms to multidrug resistance and virulence (4–9). In addition to the DBD, which is generally localized in the N-terminal part, Zn2C6-TF contain a region for activation/inhibition of the transcriptional machinery. This region, sometimes called TAD (TransActivation Domain), is present in many eukaryotic TF from yeast to humans (10) and is generally found in the C-terminal part of the proteins.

Comparative sequence analyses of the Zn2C6-TF family, initiated in the 1990s, revealed the existence of conserved regions between the Zn2C6 DBD and the TAD (11). One of these regions, named MHR (Middle Homology Region) (2), is composed of three conserved motifs involving about 80 amino acids. The MHR (also known as Fungal_trans) was extended to eight consecutive conserved motifs embedded in a large functional domain ranging from 225 to 405 residues (12), which is, like the Zn2C6 DBD, specific to fungal species and represents the second largest fungal-specific TF class (8). A mean secondary structure prediction performed on the eight motifs suggested that they are mainly composed of α-helices. Ten years later, the crystal structure of Cep3 (13, 14), one of the sequences included in the analysis (12), confirmed that the eight motifs are included in an all-alpha domain, hereafter called MHD (Middle Homology Domain). To date, Cep3 remains the only MHD-containing protein whose 3D structure has been experimentally determined.

The functional role of the MHD remains largely elusive, although it has been postulated that the fungal-specific Zn2C6-MHD TF might correspond to the metazoan nuclear receptors, with the MHD echoing the metazoan Ligand Binding Domain involved in the regulation of the TF activity (notably in an inhibitory function) and/or in the regulation and recognition of effectors (15). Furthermore, it has been postulated that the MHD might also participate in DNA target discrimination (16, 17).

In terms of protein domain organization, the domain pair or bigram (18) composed of the Zn2C6 DBD combined with the MHD is the most frequent in the Zn2C6-TF family. For example, of the 54 Zn2C6-TF from the *Saccharomyces cerevisiae* S288C strain, 44 contain a Zn2C6-MHD domain pair (12). Strikingly, in the InterPro protein family database (19), approximately one third of the proteins exhibiting an MHD (InterPro ID: IPR007219) are not predicted to contain a zinc finger motif of the Zn2C6 or C2H2 types (InterPro ID: IPR001138 or IPR013087). These TF, which apparently lack a DNA binding activity, represent the second largest fungal TF class after the Zn2C6 TF and will be called ‘MHD-only’ hereafter. Except for some rare exceptions (20), MHD-only TF have not been confirmed experimentally and there is some debate about whether the MHD can act independently. For example, in all experimentally proven TF listed in the TRANSFAC database, the MHD is always located downstream of a DBD (5).

A recent genome-wide study of the complement of TF in the fungus *Aspergillus nidulans* revealed numerous discrepancies between the predicted gene sequence annotations and the experimental transcriptomic data, with approximately 30% of the TF-coding genes needing some type of re-annotation (21). Among the badly predicted TF, a large majority (78%) concern the Zn2C6- and/or MHD-containing sequences which frequently exhibit non-predicted or non-processed introns leading to premature stop codons and erroneous annotation. It is interesting that most of the *A. nidulans* MHD-only proteins have a domain with predicted DNA binding (mainly of the Zn2C6 type) after RNA sequence analysis. These high-throughput experimental results prompted us to reassess the fungal-specific MHD-containing TF family, by developing a domain-centric approach that takes into account potential mispredictions of protein sequences.

As a starting point, we collected proteins containing an MHD from three sequence databases with different levels of human expertise involved in the genome annotation process. First, the Saccharomyces Genome Database (SGD) is dedicated to the budding yeast *S. cerevisiae* (22) and provides comprehensive information including protein sequences from a collection of 48 *S. cerevisiae* strain genomes. Second, the UniProtKB/Swiss-Prot database is the expertly curated component of UniProtKB (23). Third, the UniProtKB/TrEMBL database contains computer-generated annotations for all translations of the EMBL nucleotide sequence entries. We then focused our analysis on the MHD-only TF by applying a specific protocol using different DBD-MHD combinations to identify potentially mispredicted genes in available fungal genomic sequences, and especially mispredictions that affected the protein domain organization.

Our large-scale analysis of almost 17000 MHD-only TF allowed us to verify that at least 90% of them possess upstream genomic sequence regions coding for a DBD, mostly of the Zn2C6 type. These results suggest that the vast majority of the MHD-only TF sequences present in public databases result from annotation errors, and that the Zn2C6-MHD domain pair represents a canonical domain signature defining a new family of TF composed of two fungal-specific domains.

## RESULTS

### MHD-containing proteins in the SGD database

We first analyzed the MHD-containing proteins in the SGD, a database that provides comprehensive integrated information for *S. cerevisiae*. The reference (S288C) strain of *S. cerevisiae* contains 44 proteins with an MHD (Table S1), all of them exhibiting a Zn2C6-MHD domain pair (12). In addition to S288C, the SGD contains genome assemblies and gene annotations for a further 47 strains of *S. cerevisiae* (Table S2). For each of these 47 strains, we extracted the annotated orthologs of the 44 S288C Zn2C6-MHD proteins. If all orthologs are conserved in all strains, we would expect a total of 2068 orthologs (44 orthologs from each of the 47 strains), but only 1793 orthologous sequences were found in the SGD (Table 1). In other words, 275 (13%) orthologous sequences were not predicted. Furthermore, for the 1793 predicted sequences, 253 (14%) of them did not contain the conserved CX{2}CX{6}CX{5,16}CX{2}CX{6,8}C motif and were considered to be potentially mispredicted genes.

**Table 1.**
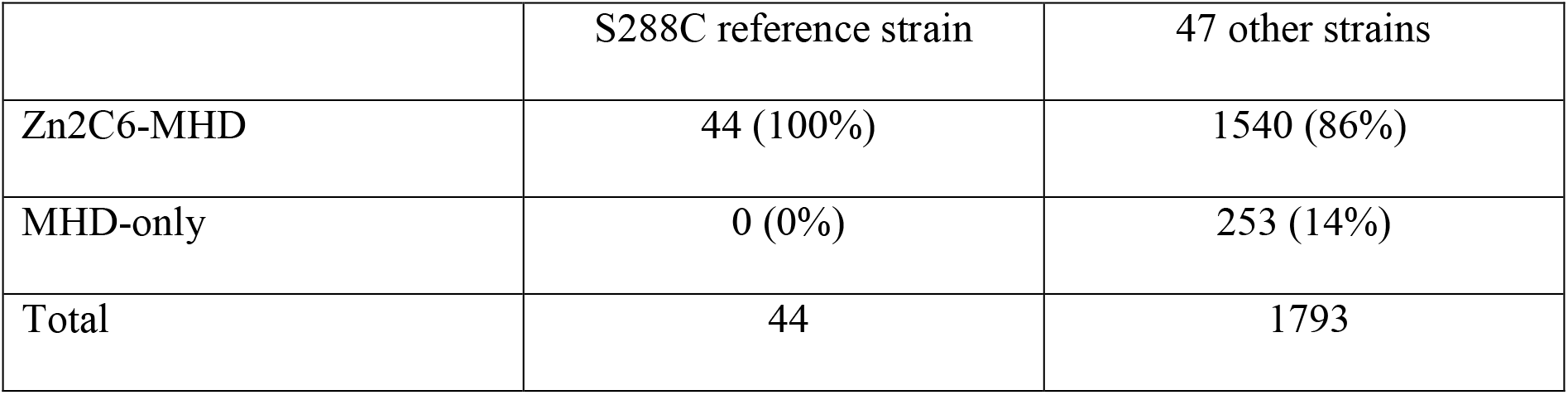
Domain annotations of MHD containing proteins in the SGD database. The numbers in brackets indicate the proportion of sequences with each domain combination, with respect to the total number of sequences.

To investigate the causes of the 253 genes with potential errors, we used the 44 S288C protein sequences to search the corresponding genome assemblies in the SGD using TBLASTN (Table 2). In the majority of cases, significant hits to genomic regions were found and the protein sequence errors could be linked to genome sequencing or assembly issues. Indeed, for 190 (75%) of the 253 proteins, the S288C protein sequence matched to multiple segments of a single genome scaffold with sequence identity of at least 95%, although the matching segments were found in different reading frames. These frameshifts were mainly due to the insertion of one or two bases in the genome sequence of the *S. cerevisiae* strain, as compared to the S288C sequence. A further nine sequences were found split over multiple scaffolds. For 46 (18%) of the 253 proteins, the S288C protein sequence matched the genome scaffold with a coverage of 100% and sequence identity of at least 95%, and we concluded that the absence of the Zn2C6 domain was due to a wrongly predicted start codon.

**Table 2.** Probable sources of protein sequence prediction errors in the SGD database. The numbers in brackets indicate the proportion of each cause of error with respect to the total number of errors detected.

| Error type | Probable cause of error | MHD-only |
| --- | --- | --- |
| Genome sequence error | Frameshift | 190 (75%) |
|  | 2 or more scaffolds | 9 (3%) |
| Gene prediction error | Wrong start codon | 46 (18%) |
| Undetermined | Undetermined | 8 (3%) |
| Total |  | 253 |

We then tried to correct the 253 erroneous sequences by reconstructing the protein sequence from the TBLASTN genome hits. For 243 (95%) sequences, a complete Zn2C6 domain could be found upstream of the MHD domain (Table S3). The full-length sequences are provided as a Fasta file. After taking into account the detected gene prediction errors, only ten of the 253 MHD-only proteins remained for which a Zn2C6 domain was not found. These included the nine gene sequences split over multiple scaffolds, which could not be resolved due to the genome assembly issues, and one sequence with a frameshift error (although manual analysis of this genome sequence indicates a frameshift error affecting one of the conserved cysteines in the Zn2C6 domain).

In summary, no convincing evidence of MHD-only proteins was found in any of the 47 *S. cerevisiae* strains analyzed here and all the identified MHD located in reliable genome sequence scaffolds were associated with upstream Zn2C6 domains.

### MHD-containing proteins in the Uniprot database

We then queried the UniProt database for all proteins annotated with the MHD (Interpro ID: IPR007219), resulting in a total of 126861 proteins, with 126691 in the unreviewed TrEMBL section and 170 in the reviewed Swiss-Prot section. The MHD containing proteins have a wide range of domain architectures, with 1905 different architectures listed in the Interpro database, although the most frequent domain pairs are: (i) MHD with a Zn2C6 DBD (IPR001138), (ii) MHD-only, and (iii) MHD with one or two C2H2 DBD (IPR013087), as shown in Figure 1 and Table S4.

**Figure 1.**
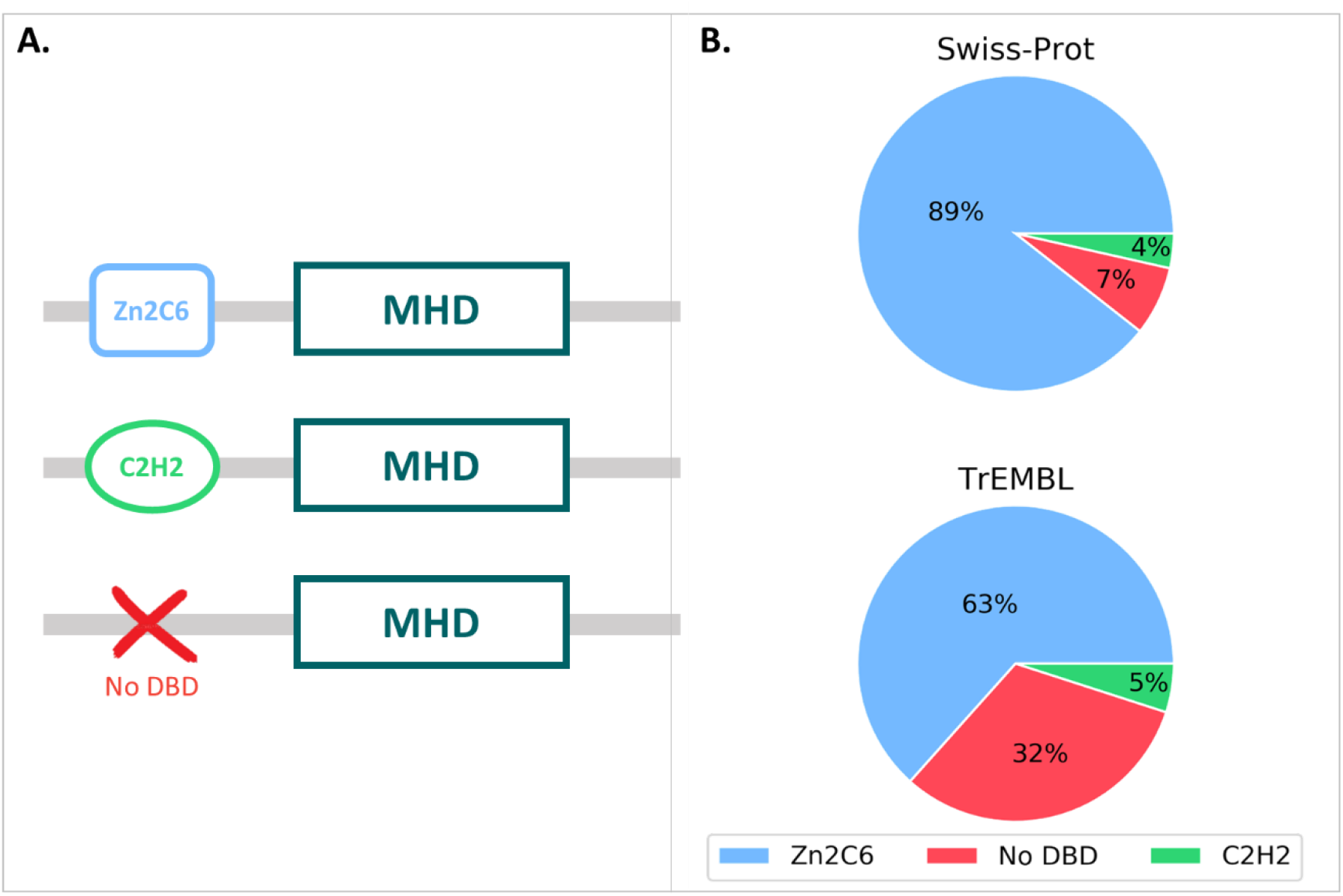
A. MHD domain pairs found in the UniProtKB. B. Proportion of each domain pair found in the Swiss-Prot and TrEMBL sections, with respect to the total number of MHD-containing sequences.

For the 170 proteins from the Swiss-Prot section, nearly 90% contain the Zn2C6-MHD domain pair, although this combination is found in a smaller proportion of the TrEmbl proteins with only 63.4%. Conversely, the proportion of MHD-only proteins lacking an annotated DBD is much higher in TrEmbl (31.6%) than in Swiss-Prot (7.1%).

### Manual analysis of the 12 Swiss-Prot MHD-only sequences

Since Swiss-Prot entries are curated by experts, we manually investigated the twelve MHD-only sequences in this database (Table S5). Where possible, we extracted the corresponding genome sequence from either ENSEMBL (24) or GENBANK (25) databases, and translated the genome sequence in the three frames to search for potential DBD encoding regions. This was not possible for four of the twelve sequences. For Q5AR44 and A0A5C1RF03, the gene is located at the start of a contig and the genome region upstream of the annotated gene is not available. For B8NJG5, according to the ENSEMBL database, the upstream gene codes for a small protein coding a Zn2C6 DBD. Finally, A6SSW6 is a shorter protein of length 179 (compared to > 500 for the other Swiss-Prot proteins), and the MHD hit in the InterPro database is a partial domain. The genome sequence (FR718884) is annotated as “possibly a relic of a transcription factor”.

For all remaining eight proteins, a potential DBD sequence is found either within the existing annotated gene *via* alternative splicing, or the proximal 5’ region (<1000 nt) *via* an alternative start codon or in a new exon (Figure 2).

**Figure 2.**
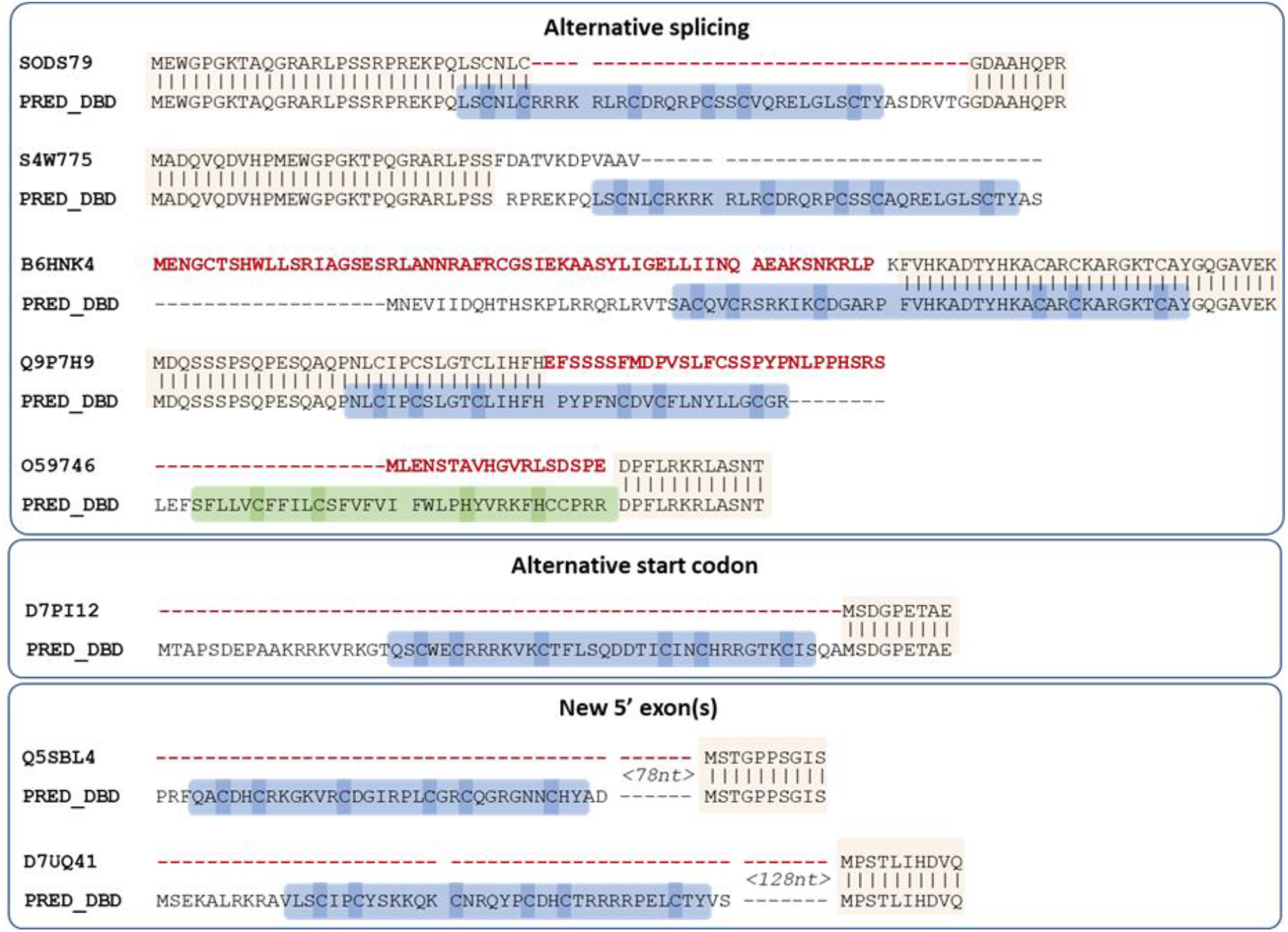
Proposed new sequences for missing DBD of Swiss-Prot MHD proteins. Conserved regions between the existing Swiss-Prot sequence and the proposed sequence are indicated by vertical lines. Regions in the existing Swiss-Prot sequence shown in red are replaced in the predicted sequence, while the predicted DBD is outlined in blue (Zn2C6) or green (C2H2), with cysteines/histidines corresponding to potential zinc binding amino acids highlighted in blue. Spaces in the sequences indicate annotated or predicted splice sites.

### Automatic analysis of 16760 TrEMBL MHD-only sequences

Based on the manual analysis of Swiss-Prot described in the previous section, an automatic protocol was developed to analyze potentially erroneous sequences retrieved from the UniProt/TrEMBL database. The first step in the protocol involved identifying the corresponding genomic sequences in the ENSEMBL database. This resulted in a set of 16760 sequences that were used as input for the main error detection step (see Methods). Two different methods were implemented to locate genomic regions within or upstream of the gene that could encode the missing DBD, using either a local or global alignment approach. Figure 3 shows the number of DBD identified by the two methods. By integrating the results of the local and global alignment searches, DBD sequences could be proposed for 14482 (86%) of the 16760 MHD-only sequences tested (Table S6). The proposed DBD sequences are provided in Supplementary File 2). Most of these sequences correspond to a Zn2C6 domain (82%), with a smaller proportion of C2H2 domains (4%), which correlates well with the proportions found in the manually curated Swiss-Prot section.

**Figure 3.**
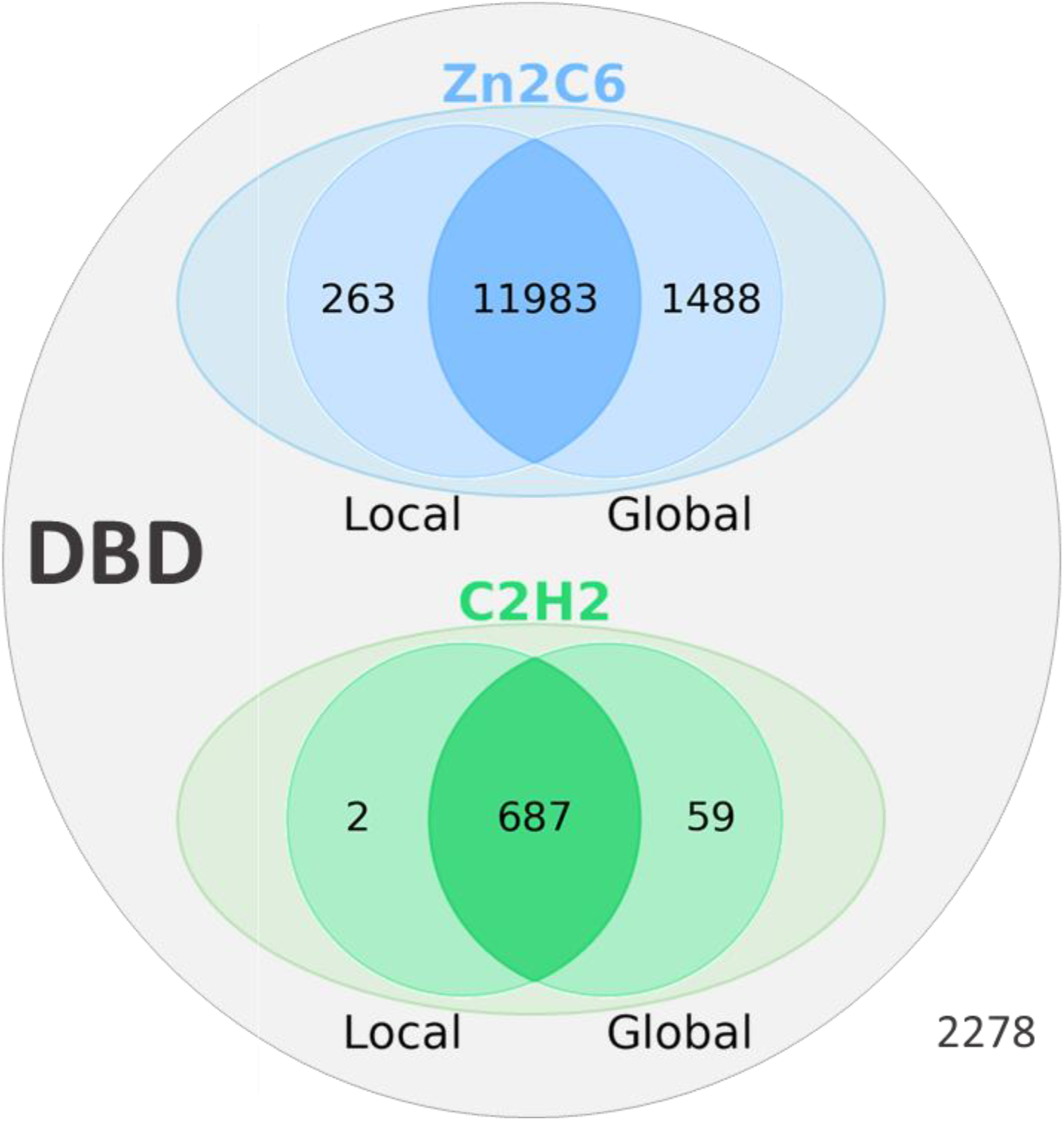
Results of the error identification step in 16760 MHD-only TF sequences from the TrEMBL database. Number of DBD (Zn2C6 or C2H2) identified by local and global alignment methods.

For the 2278 (13.6%) proteins with no DBD identified by our automatic protocol, we then investigated potential causes for the erroneous sequences. Partial hits, with hmmsearch scores below the defined threshold and part of the conserved Zn2C6 or C2H2 motifs (see Methods), were found in 905 sequences. These might indicate complex exon/intron structures that were partly mispredicted by our protocol (an example is shown in the following section) or might be caused by genome sequencing or assembly issues. For example, undefined regions in the genomic sequences, represented by ‘N’ characters, were found in 1363 of the 2278 proteins. Other reasons for not identifying a DBD include (i) the DBD is located more than 1000 nucleotides upstream of the gene, (ii) the related sequence is not conserved enough to allow protein-DNA alignment of the DBD.

### Reassessment of domain pairs in MHD-containing sequences

If the missing DBD sequences proposed here were integrated into the public databases, the number of MHD-only proteins would be reduced from 12 to 4 for Swiss-Prot, and from 16760 to 2278 for TrEMBL (Figure 4 and Table S6). More importantly perhaps, this would lead to a significant difference in the distribution of domain pairs present in MHD-containing sequences. In the public databases, this distribution is 65%, 5% and 30% for Zn2C6-MHD, C2H2-MHD and MHD-only respectively. However, our error-aware protocol indicates that the true distribution is closer to 90%, 6% and 4% for Zn2C6-MHD, C2H2-MHD and MHD-only respectively.

**Figure 4.**
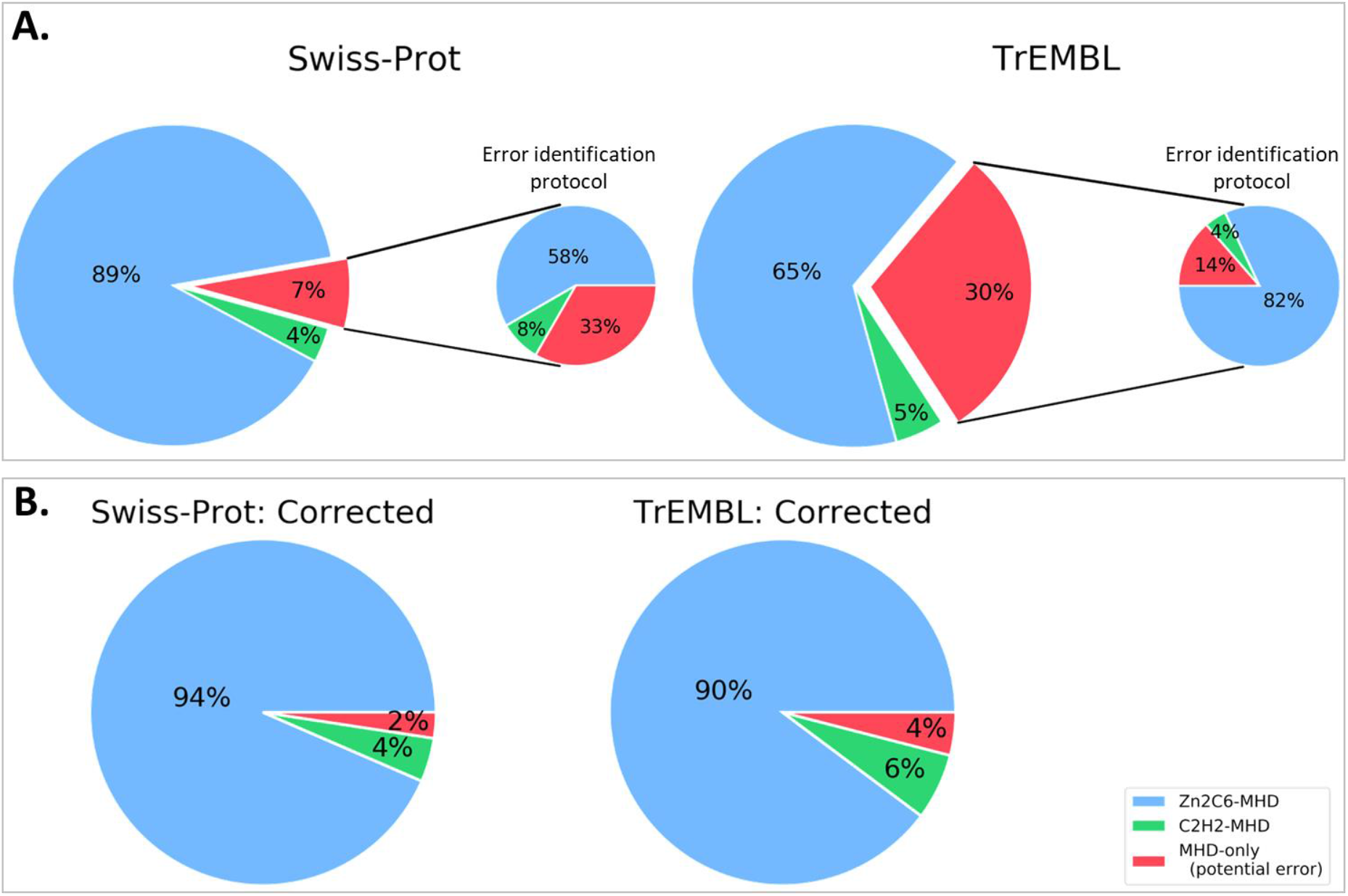
Proportion of sequences with Zn2C6-MHD (blue) or C2H2-MHD (green) domain combinations in A. public databases: Swiss-Prot and TrEMBL (sequences mapped to ENSEMBL only) and B. after applying our error identification protocol. Proportion of potentially erroneous sequences lacking a DBD is shown in red.

Interestingly, it has been shown previously that there is a significant difference in the TF repertoire of ascomycete and basidiomycete fungi (6), and in particular that the Zn2C6 family (33%) is much more prevalent than the C2H2 (10%) in ascomycete TF, compared to basidiomycete TF (20% and 15% for Zn2C6 and C2H2 respectively). Despite this overall enrichment of C2H2 in the basidiomycete TF, within the MHD sequences, the proportions of ZN2C6 and C2H2 are similar in both clades (90% and 7% for Ascomycota compared to 92% and 3% for Basidiomycota) (Table S7).

The level of gene annotation errors is of course dependent on the quality of the genome sequencing, assembly and annotation. A small number of well-characterized organisms had no MHD-only sequences in the Uniprot database, including model organisms like *S. cerevisiae*, *Yarrowia lipolytica*, or *Ustilago maydis*. Nevertheless, some model organisms had a small number of MHD-only sequences, for example *Schizosaccharomyces pombe* has 27 proteins annotated with an MHD, of which two proteins had no DBD, or *Candida albicans* with 28 MHD-containing proteins, of which only one has no DBD: Q5AJ63_CANAL (Figure 5).

**Figure 5.**
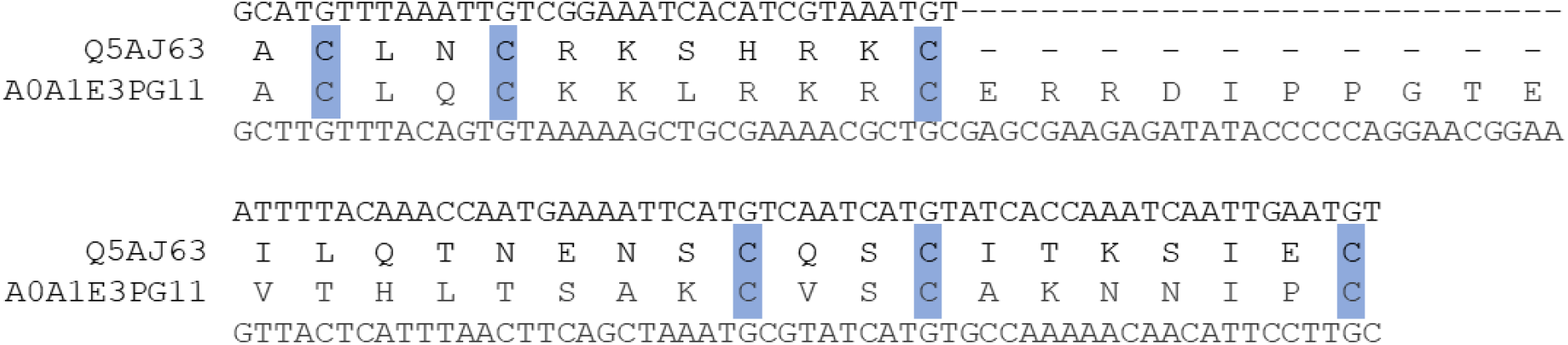
Predicted DBD for Candida albicans sequence Q5AJ63_CANAL, aligned with neighbor sequence A0A1E3PG11_9ASCO (hmmsearch E-value = 6.5e-10). Conserved cysteines characterizing the Zn2C6 DBD are highlighted in blue.

At the other extreme, *Rhizopus delemar* has 34 MHD-containing proteins of which only 6 are also annotated with a DBD, i.e. 82% are MHD-only proteins. According to our protocol, DBD could be detected for a further 24 MHD proteins and only 4 (12%) lack a DBD. Further manual analysis of these 4 MHD-only proteins showed that two genes (I1BR31, I1C782) have regions coding complete Zn2C6 domains within 1500 nt upstream of the 5’ end, while one gene (I1CF81) has a partial hit within the default threshold of 1000 nt upstream of the 5’ end (−490 to −192). According to the public databases, I1CF81 is coded by a gene with one exon and contains one known domain, the MHD. The partial DBD hit for I1CF81 in fact contains a misprediction of a short exon coding for 2 amino acids, as shown in Figure 6.

**Figure 6.**
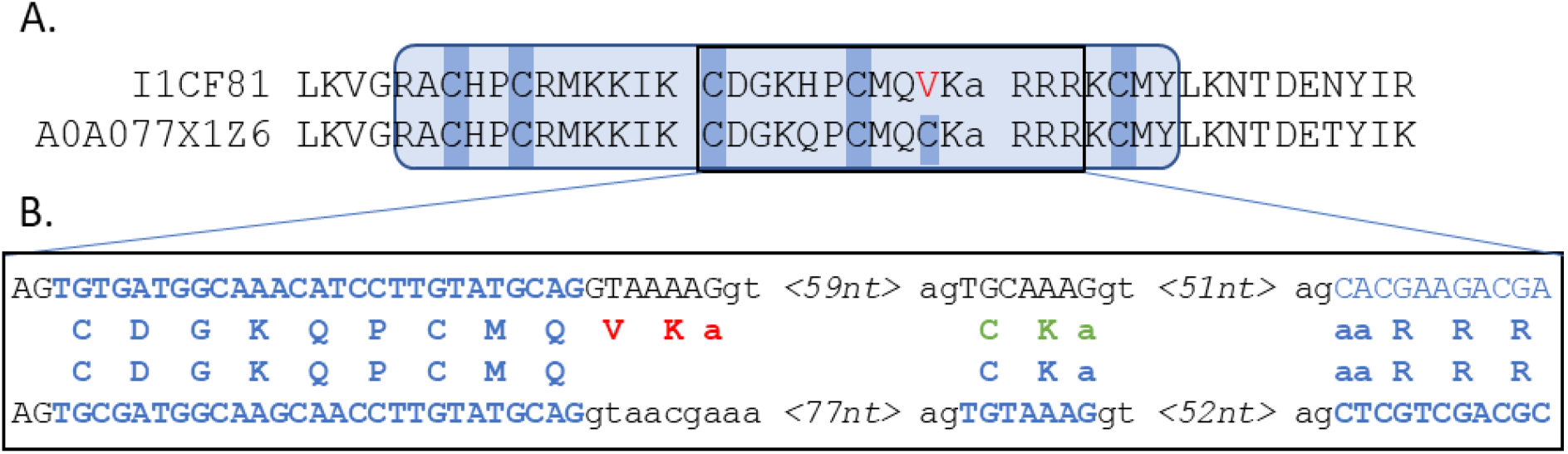
A. Protein alignment of query I1CF81 with the neighbor A0A077X1Z6, showing the partial hit identified by the protocol, where the predicted sequence presents five of the six conserved cysteines that characterize the Zn2C6 DBD (hmmsearch E-value = 1.5e-11). Exon/intron boundaries are indicated by gaps in the sequences. B. Genome-level comparison of query I1CF81 with the neighbor A0A077X1Z6, showing correctly predicted amino acids (blue), protocol mispredicted amino acids (red) and alternative manual prediction (green).

## DISCUSSION

In this work, we reassessed the domain composition of the fungal MHD-only TF, which represent almost one third of the proteins exhibiting an MHD in the public databases. We focused on protein domains because these are the basic functional, evolutionary, and structural units that shape proteins. Domains can function independently in single-domain proteins or cooperatively in multi-domain proteins. Thus, some domains always co-occur with specific functional partners, whereas others are more versatile (26). After the Zn2C6-TF family, the MHD-only TF constitute the second most abundant fungal-specific TF family, and it remains to be determined whether the MHD can effectively act independently (5, 8).

In order to characterize the full complement of MHD-only sequences in public databases, we exploited the fact that TF are generally composed of various domain pairs that function cooperatively. The concept of associated domains in proteins has also been called ‘supra-domains’ (27), DASSEM units (28), or domain co-occurrence (29, 30). It has been used to improve protein domain identification, protein function prediction and homology/evolutionary analyses. For example, domain ‘context’ is used to improve protein domain prediction in the Pfam protein family database (31).

To assess the domain organization of TF in public databases, we defined a three-level approach with increasing complexity starting from the analysis of MHD-only sequences in 48 *S. cerevisiae* strains available in SGD, followed by the analysis of the sequences present in the expert curated Swiss-Prot database and finally, in the automatic computer-generated TrEMBL database. In the reference *S. cerevisiae* S288C strain, no MHD-only TF are present and we showed that, for 95% of the MHD-only TF observed in the other *S. cerevisiae* strains, a complete Zn2C6 domain is located upstream of the MHD domain. These discrepancies in the domain annotations of the proteins were generally linked to either genome sequence or gene prediction errors. Our analysis highlights the unexpectedly high rate of sequence errors in these very closely related genomes. This high error rate was confirmed by our analysis of the 12 MHD-only proteins present in the expert curated Swiss-Prot database, since a DBD could be identified for all the proteins whose corresponding genomic sequence was available. Finally, concerning the TrEMBL proteins, our error-aware protocol showed that 89% of the MHD-only TF exhibit upstream genomic sequence regions coding for a DBD. These results are in line with the error rate of 66% observed in the study of Zn2C6-TF in *Aspergillus flavus* (32), involving manual analysis of the genomic region upstream the MHD-only proteins and emphasizing the urgent need for a systematic reassessment of these transcription-related factors lacking a DNA-binding activity. The high rate of wrongly predicted TF sequences in our study (at least 82%) is particularly surprising given that (i) fungal genome sequences are generally of better quality with fewer genome assembly errors, thanks to their relatively small, compact genomes and the low level of repetitive sequences in most fungi (33), (ii) fungi serve as model eukaryotic organisms and a wide range of diverse genomes have been sequenced and annotated (SGD, Génolevures: genolevures.org, 1000 Fungal Genomes Project: mycocosm.jgi.doe.gov), (iii) gene annotation in the fungi is facilitated by the relatively streamlined gene structures and transcriptional processes in these organisms with few and typically short introns rarely implicated in alternative splicing. Our results clearly indicate that all these fungal features, which should improve the gene prediction quality, do not limit the error rates at least in the studied TF family.

This notion of errors in public protein databases is a recurrent problem (34–36) and substantial efforts have been invested to identify and correct gene annotation errors (37–39). Some important causes of erroneous sequences have been identified, including the genome sequence quality and gene structure complexity (40), as well as redundant or conflicting information in different resources or in the literature (34, 41). Consequently, it has been estimated that 40 to 60% of the protein sequences in public databases are erroneous (42–44). Typical errors include missing exons, non-coding sequence retention in exons, wrong exon and gene boundaries, fragmenting genes and merging neighboring genes. Thus, the development of automated methods to identify and correct mispredicted protein sequences remains an important research topic (43, 45–47). Our study showing high error rates at a family-wide level is further evidence of the potential of domain-centric approaches for sequence annotation correction and of the urgent need to clean the databases.

At the functional level, our study shows that MHD-only TF sequences result predominantly from annotation errors, and are not widespread in fungi as previously described. As Zn2C6-TF function mainly as homodimers or heterodimers, this implies that the number of sequence-specific TF and the array of control DNA-sequences in target genes need to be reconsidered, as well as the degree of combinatorial regulation involved in the wide range of fungal processes controlled by this TF family (4–9). The results presented here represent important evidence that the fungal-specific MHD should be considered as part of a synergistic domain pair with a zinc finger DBD, largely composed of the fungal-specific Zn2C6 type. As a consequence, the Zn2C6-MHD architecture defines the most widely distributed and abundant TF family specific to fungi. We propose to name this family CeGAL after its most characterized members: **Ce**p3, whose 3D structure has been determined, and **GAL**4, the archetypal eukaryotic TF. This definition should not only improve the quality of future fungal genome annotations, but should provide better discrimination of the so-called GAL4-like regulators defined according to the presence of a Zn2C6 domain and which include TF with diverse domain architectures.

More importantly, this will also contribute to a better understanding of fungal Gene Regulatory Networks (GRN), that aim to define the complete set of regulatory interactions between TFs and their target genes at a species level. Classically, GRN analyses combine an initial step of genome-wide characterization of TF families with experimental data related to transcriptional effects of TF deletion/overexpression, chromatin immunoprecipitation (ChIP)-based TF binding data, protein-protein interactions or pairs of genes involved in genetic interactions (48). However, as GRN studies generally exclude proteins lacking a DBD, the complete repertoire of fungal TF is frequently underestimated. It will be essential that future comprehensive, genome-wide TF studies consider the CeGAL DBD-MHD family as a synergistic domain pair of DBD and MHD, in order to provide new and unbiased insights for regulatory network analysis.

## MATERIALS AND METHODS

### Collection of SGD sequences

The 44 proteins from the *S. cerevisiae* S288C strain with a Zn2C6-MHD domain organization (12) were identified in the SGD database (Table S1), and their annotated orthologs in the 47 available strains (Table S2) were downloaded from the SGD web site (http://sgd-archive.yeastgenome.org/sequence/strains/strain_alignments.tar). Genome assemblies for the 47 strains were also downloaded from the SGD web site (http://sgd-archive.yeastgenome.org/sequence/strains). Ortholog sequences that did not contain the conserved CX{2}CX{6}CX{5,16}CX{2}CX{6,8}C motif were considered to be potentially erroneous.

For each potentially erroneous sequence, we performed a TBLASTN alignment of the S288C reference protein sequence with the corresponding genome assembly. We then tried to identify the causes of the erroneous sequences. First, if TBLASTN hits were found on multiple scaffolds (with percent identity >95% and length>20 amino acids), we assumed that the misprediction was due to a genome assembly issue. If multiple TBLASTN hits (with percent identity >95% and length>20 amino acids) were found on a single scaffold, but in different reading frames, we assumed that the misprediction was due to a sequence insertion leading to a frameshift error. If a TBLASTN hit was found with percent identity >95% and coverage =100%, we assumed that the misprediction was due to a wrongly predicted start codon.

In order to propose a corrected sequence, a protein sequence was then reconstructed from the TBLASTN hits found on the same scaffold. This corrected protein sequence was searched for the conserved CX{2}CX{6}CX{5,16}CX{2}CX{6,8}C motif.

### Collection of UniprotKB sequences

Fungal proteins were identified in the UniProt 2022_01 database (23), by querying for proteins annotated with the Interpro entry IPR007219: *Transcription_factor_dom_fun* or *Fungal_trans*, which covers the Middle Homology Domain (MHD) specific to these transcription factors. Domain architectures of all proteins containing an MHD were then extracted from the Interpro v86.0 database (19). The 37648 UniProt sequences annotated with an MHD, but no DBD, were considered to be potentially erroneous (MHD-only). Potentially erroneous sequences from reviewed UniProt/Swiss-Prot entries and unreviewed UniProt/TrEmbl entries were processed separately: the 12 UniProt/Swiss-Prot proteins with no DBD were analyzed manually, while the 37634 UniProt/TrEMBL sequences were input to the error identification protocol, outlined in the schema in Figure 7, and described in detail in the following sections.

**Figure 7.**
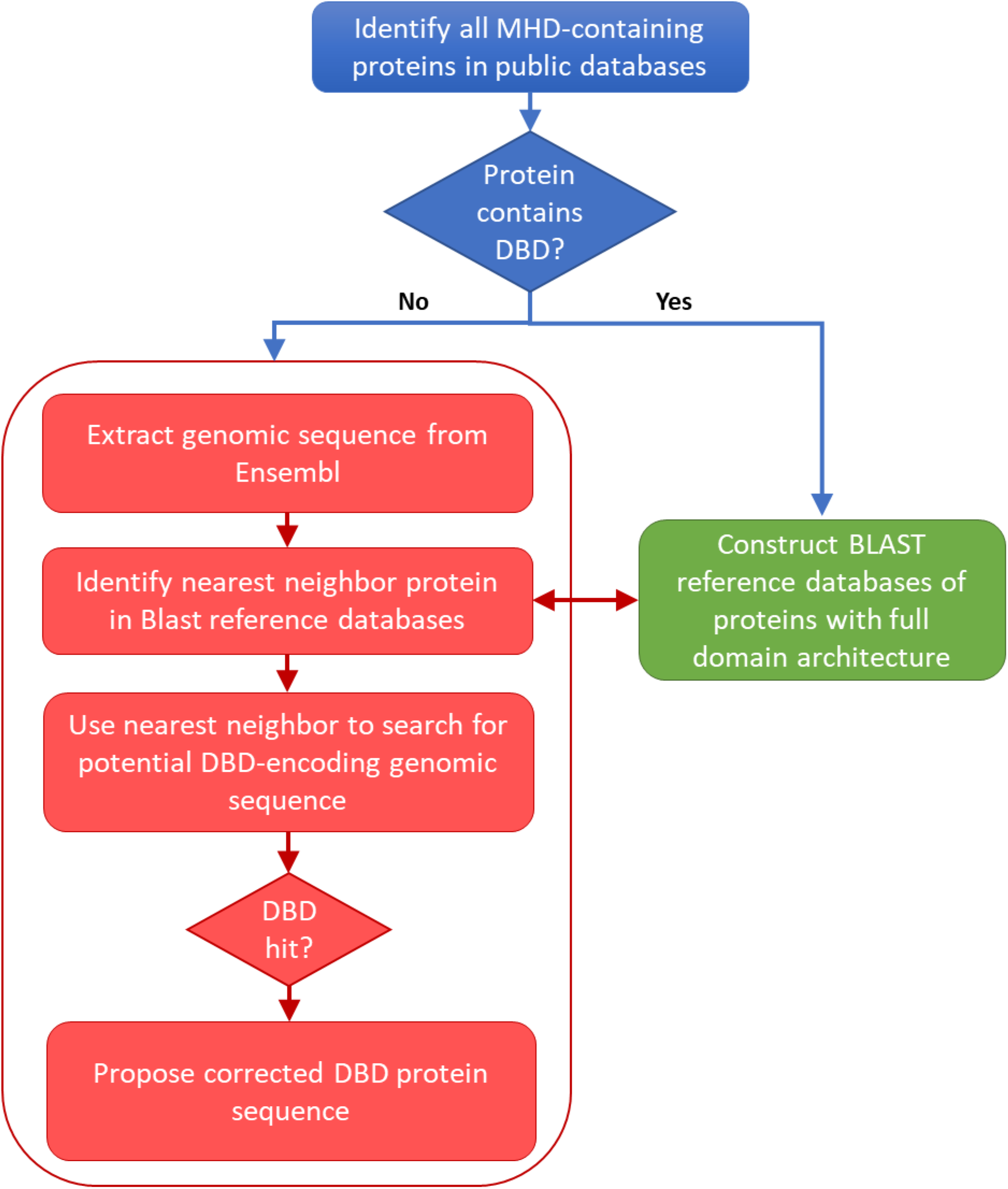
Schema of the protocol used to locate potential errors in sequences retrieved from a public database.

### Construction of BLAST databases of proteins with full domain architecture

UniProt sequences containing an MHD (IPR007219) in combination with a DBD were used to construct BLAST reference databases. Two BLAST databases were constructed: one for each of the two main DBD types, namely Zn2C6 (IPR001138) and C2H2 (IPR013087), found in this TF family to which the well-studied GAL4 protein belongs. The Zn2C6 BLAST reference database contains 80456 sequences, while the C2H2 BLAST reference database contains 6314 sequences.

### Extraction of genomic sequences

For all potentially erroneous sequences in UniProt, the corresponding genomic sequences were extracted from the Ensembl database (24), when an Ensembl cross-reference was available in the UniProt database. To improve detection of missing DBD, the full length gene sequences were retrieved together with an additional 1000 nucleotides upstream of the 5’ end of the gene. For the 37634 potential error sequences, 16760 DNA sequences were found in the Ensembl database.

### Identification of nearest neighbor reference sequences

For each MHD-only sequence, BLASTP searches were performed in the two BLAST reference databases containing Zn2C6 and C2H2 sequences in combination with an MHD. The nearest neighbor sequences with the required domain combination (i.e. DBD and MHD) were selected, if a BLASTP hit was identified with E-value < 0.005. Figure 8 shows the E-value distribution of BLASTP hits obtained for all neighbor sequence searches.

**Figure 8.**
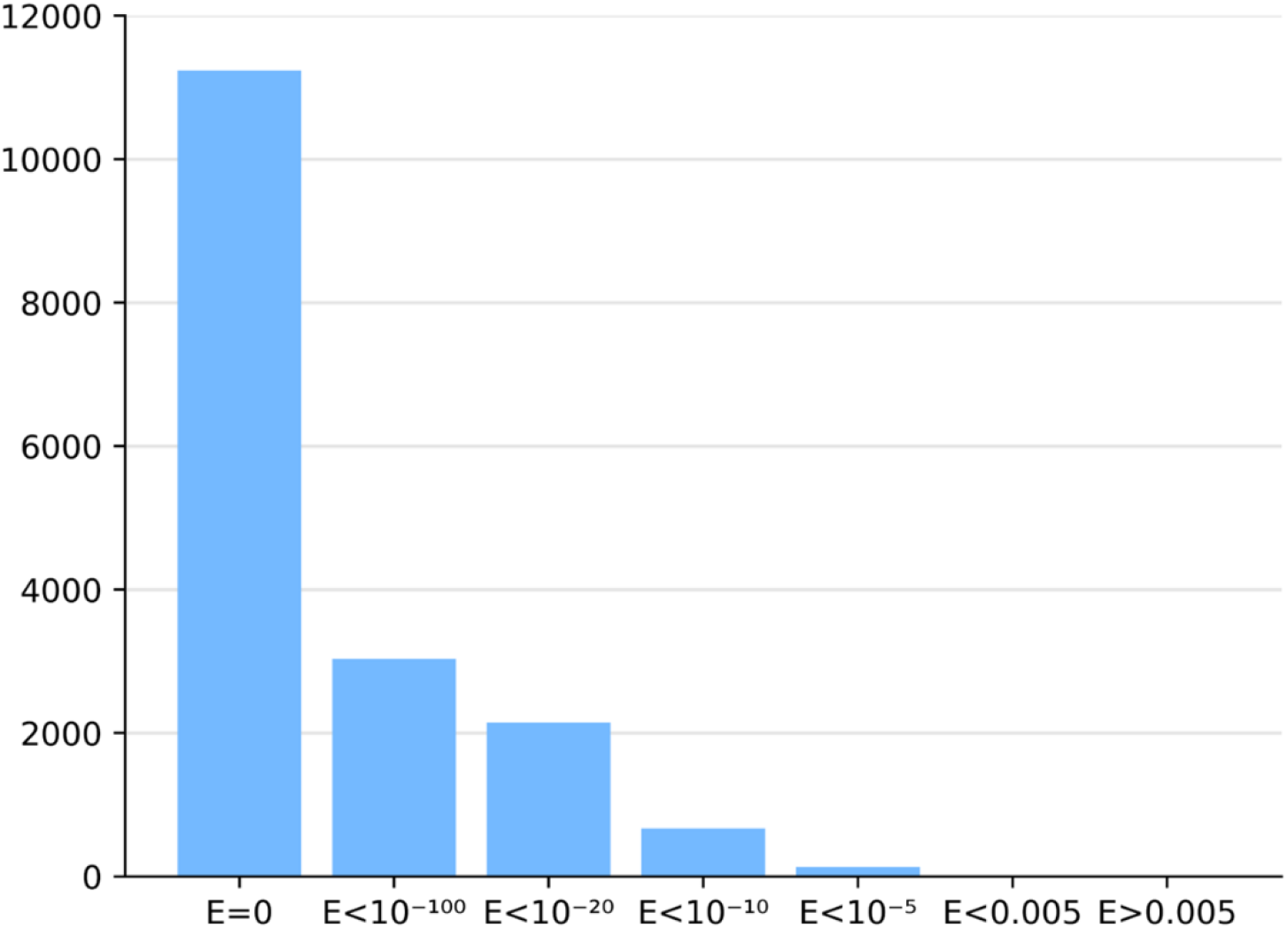
Histogram of BLASTP E-values in neighbor identification step. Three neighbor sequences were identified with E-value> =0.005 and were removed from the following steps.

### Identification of missing DBD sequences

For each MHD-only sequence with a BLASTP hit to a nearest neighbor reference protein, two complementary approaches were implemented to search for the missing DBD sequence. First, a local TBLASTN search was performed in the genomic sequence of the potential error sequence, using the protein DBD sequence segment of the nearest neighbor as a query. TBLASTN alignments with E-value < 0.0001 were taken into account. Second, a global pairwise alignment was performed between the genomic sequence of the MHD-only sequence and the full-length protein sequence of the nearest neighbor, using the Prosplign software developed by the NCBI (https://www.ncbi.nlm.nih.gov/sutils/static/prosplign/prosplign.html). Pairwise alignments obtained from TBLASTN and Prosplign were analyzed to identify potential DBD-encoding sequence segments in the erroneous sequences. Finally, the potential DBD-encoding sequence segments were compared to an HMM representing the DBD downloaded from the Pfam protein family database (31): PF00172 for the Zn2C6 DBD and PF00096 for the C2H2 DBD. To do this, the hhmsearch program from the HMMER suite (49) was used and sequences with an E-value < 0.1 were considered as hits. In addition to the hmmsearch E-value and to eliminate a number of false positive hits, we also checked for conserved amino acids: for Zn2C6 DBD, two occurrences of the pattern C-x(2)-C (where x is any amino acid) were required, while for C2H2 DBD, one occurrence for each of the C-x(2,4)- C and H-x(3,5)-H patterns were required.

In order to determine whether our sequence curation protocol over-predicts DBDs in the potentially erroneous sequences, we used the same error identification protocol to search for Zn2C6 DBD in the C-terminal region of the proteins (i.e. downstream of the MHD). The results of this analysis are described in the Supplementary methods).

## DATA AVAILABILITY

The genomic and protein sequences supporting the conclusions of this article are available in public databases: SGD, UniprotKB, Ensembl and GENBANK. The full-length sequences for the corrected MHD-containing proteins from the SGD database, and the corrected sequences for the missing DBD in the MHD-containing proteins from the TrEMBL database are available at http://git.lbgi.fr/julie/CeGAL-DBD.

## ACKNOWLEDGEMENTS

We thank the members of the BiGEst bioinformatics platform for their assistance. This work was supported by French Infrastructure Institut Français de Bioinformatique (IFB) ANR-11-INBS-0013, and Institute funds from the French Centre National de la Recherche Scientifique, the University of Strasbourg. Funding for open access charge: University of Strasbourg.

